# Knee and Hip Joint Dynamics Differ between Sprinting and Nordic Hamstring Exercises

**DOI:** 10.1101/2025.08.15.670579

**Authors:** Kristen Steudel, Nicos Haralabidis, Reed Gurchiek, Jennifer Hicks, Scott Delp

**Author notes:** Corresponding author’s (K. Steudel). These authors contributed equally to this work (Kristen Steudel, Nicos Haralabidis).

## Abstract

**Background:** Sprinting and Nordic hamstring exercise (NHE) programs are common training modalities used to reduce hamstring injury risk, but the differences in the biomechanical demands of sprinting and the NHE are unclear. The purpose of this study was to compare knee and hip joint kinematics and kinetics, and hamstrings muscle-tendon unit (MTU) length and velocity during the flight phase of sprinting and the NHE.

**Methods:** We collected motion capture and force data from fourteen young athletic participants (8 males and 6 females) as they ran at a range of speeds (4–8 m/s) and performed the NHE. We used this experimental data and a musculoskeletal model to compute joint angles, moments, work, and power and to estimate the hamstrings MTU length and velocity for all running speeds and the NHE.

**Results:** The peak knee flexion moment at running speeds of 6 m/s and above was greater than for the NHE (*p* < 0.001). Peak negative knee flexion power at all running speeds was higher than during the NHE (*p* < 0.001). Negative knee flexion work at running speeds of 6 m/s and slower was less than during the NHE (*p* < 0.001). Peak hamstrings length and lengthening velocity were greater (*p* < 0.001) for all running speeds compared to the NHE.

**Conclusion:** Sprinting puts the hamstrings at longer hamstrings lengths and higher hamstrings lengthening velocities than the NHE. The NHE requires participants to generate peak knee flexion moments that are smaller than the peak knee flexion moments generated during top speed sprinting and peak negative knee flexion powers that are less than 5% of sprinting. However, the duration of each NHE repetition is approximately 60 times longer than the hamstrings lengthening portion of the flight phase of running, resulting in comparable negative knee work. The results of this study provide necessary quantitative information to compare the biomechanical demands of sprinting and the NHE.

## 1. Introduction

Hamstring strain injuries are one of the most common injuries in sports that feature high-speed running, such as soccer,^1,2^ Australian football,^3^ and sprinting events.^4^ These injuries predominantly affect the biceps femoris long head.^5^ With an average of 18 to 22 days lost from training and competition,^6^ hamstring strain injuries impair individual and team performance.^7^ Hamstring injuries may also reduce the injured athlete’s performance after return to play^8^ and have a 13% recurrence rate.^8^ Consequently, many studies have been conducted to identify the mechanisms that lead to hamstring strain injuries and develop strategies to prevent them.

Hamstring injuries frequently occur during the swing phase of running^9^ when hamstrings reach peak force and length, perform negative work, and are highly activated.^10–13^ Hamstring training programs, including sprinting and the Nordic hamstring exercise (NHE), aim to improve the muscle’s ability to withstand these mechanical demands by stimulating hypertrophy, fascicle lengthening, and the serial addition of sarcomeres.^14–16^ NHE programs have been found to reduce hamstring strain injuries,^17,18^ increase biceps femoris long head fascicle length,^19–21^ and increase hamstrings strength.^19–21^ Programs featuring high-speed running have also been shown to prevent injuries,^22^ improve hamstrings strength^20,23^ and increase the biceps femoris long head fascicle length.^20^ A single study that compared high-speed running and the NHE showed that the NHE activates the hamstrings to high levels at joint angles similar to those at which peak hamstring activation occurs during sprinting.^24^ However, the differences in mechanical loading between the NHE and high-speed running have not been fully characterized.

The mechanics of the NHE and high-speed running differ. During a NHE repetition, the knee extends at approximately 15 °/s while hip flexion remains nearly constant.^19,21^ In contrast, during the swing phase of high-speed running, the peak knee extension and hip flexion velocity are over 800 °/s.^25^ While researchers in separate studies have used musculoskeletal modeling to quantify joint angles, joint moments, biceps femoris long head MTU length and muscle forces for the NHE^19,26,27^ and high-speed running,^10–12^ comparing the results of these studies is challenging due to the differences in musculoskeletal modeling approaches and participants. Therefore, there is a need to directly compare the mechanical loads from the NHE and high-speed running using the same musculoskeletal modeling methods and the same participants.

The purpose of our study was thus to compare the biomechanical demands of running across a range of speeds to those of the NHE. We collected motion capture data for participants performing the NHE and running at speeds between 4 and 8 m/s and used musculoskeletal modeling to quantify biomechanical demands. We examined the knee and hip flexion-extension angles, and biceps femoris long head MTU length and lengthening velocity. We also computed knee and hip flexion-extension moments, powers, and work for both training modalities to allow for a more comprehensive comparison than has previously been possible.

## 2. Methods

### 2.1. Participants

Fourteen athletes (8 males and 6 females; age: 26 ± 5 years; mass: 71.3 ± 11.1 kg; height: 1.75 ± 0.08 m) volunteered to participate in this study. Participants reported a history of regular participation in sporting activities (e.g., football, track and field, and soccer). Written informed consent was obtained from the participants, and ethical approval for the study protocol was granted by the Stanford University Institutional Review Board (IRB-67713).

### 2.2. Experimental data collection

For the first testing session, each participant performed three maximal-effort 60 m sprints at an outdoor athletics track to establish their top speed. Prior to the sprints, participants completed a self-led warm up, and they rested at least 5 minutes between sprints. Participants self-initiated the sprints from a stationary starting position. We used timing gates (Dashr Timing System, NE, USA) to measure 5 m splits between 35-40, 40-45, and 45-50 m. We calculated each participant’s top speed as the best single split from across the three sprints. To qualify for the study, 95% of the participant’s top speed had to be ≥ 7 m/s if female and ≥ 7.5 m/s if male. Seventeen participants were recruited for the study. Fourteen participants met the speed threshold with an average top speed of 8.14 ± 0.50 m/s. Participants who met the speed threshold completed a self-supervised NHE training program for at least four weeks (two sets of six repetitions twice per week) preceding biomechanics evaluation in the laboratory, regardless of whether they had prior NHE training experience.

For the laboratory testing session, participants ran at a range of speeds and performed five NHE repetitions. Participants completed their own warm-up and, during testing, rested between the running trials or NHE repetitions until they felt ready to continue. The first running speed was 4 m/s. The speed was increased by 1 m/s in each subsequent trial up to 7 m/s and then by 0.5 m/s until the speed exceeded 95% of each participant’s top speed. For example, a participant with a top speed of 8.1 m/s would have run at 4, 5, 6, 7, and 7.5 m/s. The speed cut-off was designed to mitigate the risk of injury during the study. Two participants ran at speeds up to 7 m/s, seven participants ran at speeds up to 7.5 m/s, three participants ran at speeds up to 8 m/s, and two participants ran at 4, 5, 6, 7, and 8 m/s.

We measured ground reaction forces at 2000 Hz while participants ran once at each speed on an instrumented treadmill (Bertec Corp., Columbus, OH, USA). Participants wore a safety harness that was suspended from ceiling beams. The cord attached to the harness and ceiling beams was sufficiently slack to prevent interfering with the participants’ natural running mechanics. Participants started each running trial while the treadmill was stationary, and for each trial the treadmill was accelerated and decelerated at 1 m/s^2^. Once the treadmill had accelerated to the desired speed, participants completed a minimum of 10 strides.

After the running trials, participants performed the NHE repetitions on a pair of custom-made wooden boards that measured ground reaction forces separately from each lower-limb. For each NHE repetition, participants kneeled on the boards while their ankles were each secured to one of the boards with a velcro strap. The boards were each weighed down using two 20 kg weightlifting plates. The force plates were zeroed prior to recording the trials with the weights and custom-made boards on top. Two uniaxial load cells collected ankle strap forces (HT Sensor Technology, Shaanxi, China), and two in-ground force plates (Bertec Corp., Columbus, OH, USA) collected ground reaction forces, both at 2000 Hz. The participants were instructed to complete each NHE repetition as slowly as possible^28,29^ and to lower their bodies until they could no longer hold themselves up.

We captured marker trajectories at 200 Hz using a 27 camera optical motion capture system (2 model Raptor E, 4 model Raptor 12, 7 model Kestrel 1300, 10 model Kestral 2200, and 4 model Kestral 4200, Motion Analysis, Cortex, CA, USA). Eleven cameras were positioned to capture the treadmill volume, and 16 cameras captured the NHE volume. We attached 52 retro-reflective markers to the surface of each participant’s skin and shoes using double-sided adhesive and medical tapes at palpable anatomical locations. The markers were attached bilaterally on the anterior superior iliac spine, posterior superior iliac spine, iliac crest, lateral and medial femoral epicondyles, lateral and medial malleoli, calcaneus, second and fifth metatarsal heads, acromion, lateral and medial humeral epicondyles, lateral and medial styloid processes, and unilaterally on the sternum and C7, for a total of 32 anatomical markers. An additional set of 20 tracking markers were attached to the thighs, shanks, upper arms, and lower arms. We detached the medial femoral epicondyle and malleoli markers prior to the NHE repetitions.

### 2.3. Data processing

We trimmed sprint trials to include 10 strides at the set treadmill speed. We input cleaned marker trajectories for all trials (sprints and NHE) to AddBiomechanics^30^ (Stanford University, CA, USA) to scale a generic model^31^ and calculate inverse kinematics. One repetition of the NHE was removed for one participant due to kinematic errors that resulted from marker obstruction. We low-pass filtered the kinematics and ground reaction forces using a 4th order Butterworth filter at 10 and 4 Hz for the running and NHE trials, respectively. We used the OpenSim API (version 4.4; Stanford University, CA, USA)^32–34^ in MATLAB (2024b; MathWorks Inc., Natick, MA, USA) to perform inverse dynamics and extract the biceps femoris long head MTU length and lengthening velocity. For the NHE repetitions, we applied the measured ground reaction forces for each limb to the tibia bodies.

For each steady-state running speed, we analyzed the dynamics of the first three flight phases ending with a left leg touchdown. We defined the right foot take-off as the instant when the filtered vertical ground reaction force (50 Hz cut-off; 4th order low-pass Butterworth filter) fell below 40 N and left foot touchdown when the force exceeded 40 N. Only the flight phase of sprinting was analyzed because peak biceps femoris long head MTU stretch occurs in the late flight phase,^10,12^ and the majority of the total negative work performed by the hamstrings occurs during the flight phase.^10,12^ We analyzed the dynamics of the left leg during the middle three NHE repetitions (the first and last were excluded). An NHE repetition was defined to begin when either of the filtered ankle strap forces (4 Hz cut-off; 4th order low-pass Butterworth filter) exceeded 50 N. The end of an NHE repetition was defined as the time point when the vertical velocity of either medial styloid marker first crossed zero.^26^ If neither medial styloid marker had a vertical velocity that crossed zero, the end was defined as the time point when the vertical velocity of either marker was closest to zero, following the peak downward vertical velocity.

We chose to analyze the mechanical loads for both exercises only for the portion of the movement where the hamstrings were lengthening. We used the biceps femoris long head MTU lengthening velocity as a proxy for hamstrings lengthening and chose the regions for which the velocity was positive. The mechanical load parameters compared between exercises included peak biceps femoris long head MTU length (hamstrings length), lengthening velocity (hamstrings velocity), peak knee flexion and hip extension moments, peak negative knee flexion and hip extension powers, and net knee flexion and hip extension work. To calculate knee and hip flexion-extension powers, we calculated the product of the corresponding joint angular velocity and joint moment. We integrated the knee and hip flexion-extension powers with respect to time over the identified region of hamstrings lengthening to calculate the net work. To account for size differences between participants, we normalized hamstrings length by the muscle-tendon unit length of the biceps femoris long head, with all lower extremity angles set to zero for each participant.^35^ We computed hamstrings lengthening velocities in normalized lengths per second. Moments, powers, and work were normalized to body mass.^36,37^

### 2.4. Statistical analysis

We performed all statistical analyses using RStudio (version 4.4.2).^38^ For each outcome variable (peak hamstrings length, peak hamstrings lengthening velocity, peak knee flexion moment, peak negative knee flexion power, peak hip extension moment, peak negative hip extension power, net knee flexion work, and net hip extension work), we fit a linear mixed effects model to the running speeds using the lme4 package.^39^ We calculated the sample mean of the NHE for each outcome variable. We used the emmeans package^40^ to compare the estimated marginal means from each speed in the linear mixed effects model to the NHE sample mean using a t-test with a Bonferroni correction. We used pairwise t-tests to compare the estimated marginal means at each of the whole running speeds using the contrast function from the emmeans package.

## 3. Results

During the NHE, the knee slowly extended, and the hip slowly flexed throughout each repetition, with a pronounced increase in the rate of knee extension over approximately the last 10% of the repetition (Figure 1A and B). For all running speeds, the knee extended from approximately 0-75% of the flight phase and then flexed, while the hip initially flexed and then extended. The duration of hamstrings lengthening for an NHE repetition was 4.14 ± 2.04 s, while for running, the duration of lengthening was 0.08 ± 0.01 s at 4 m/s and 0.06 ± 0.01 s at 8 m/s.

**Figure 1.**
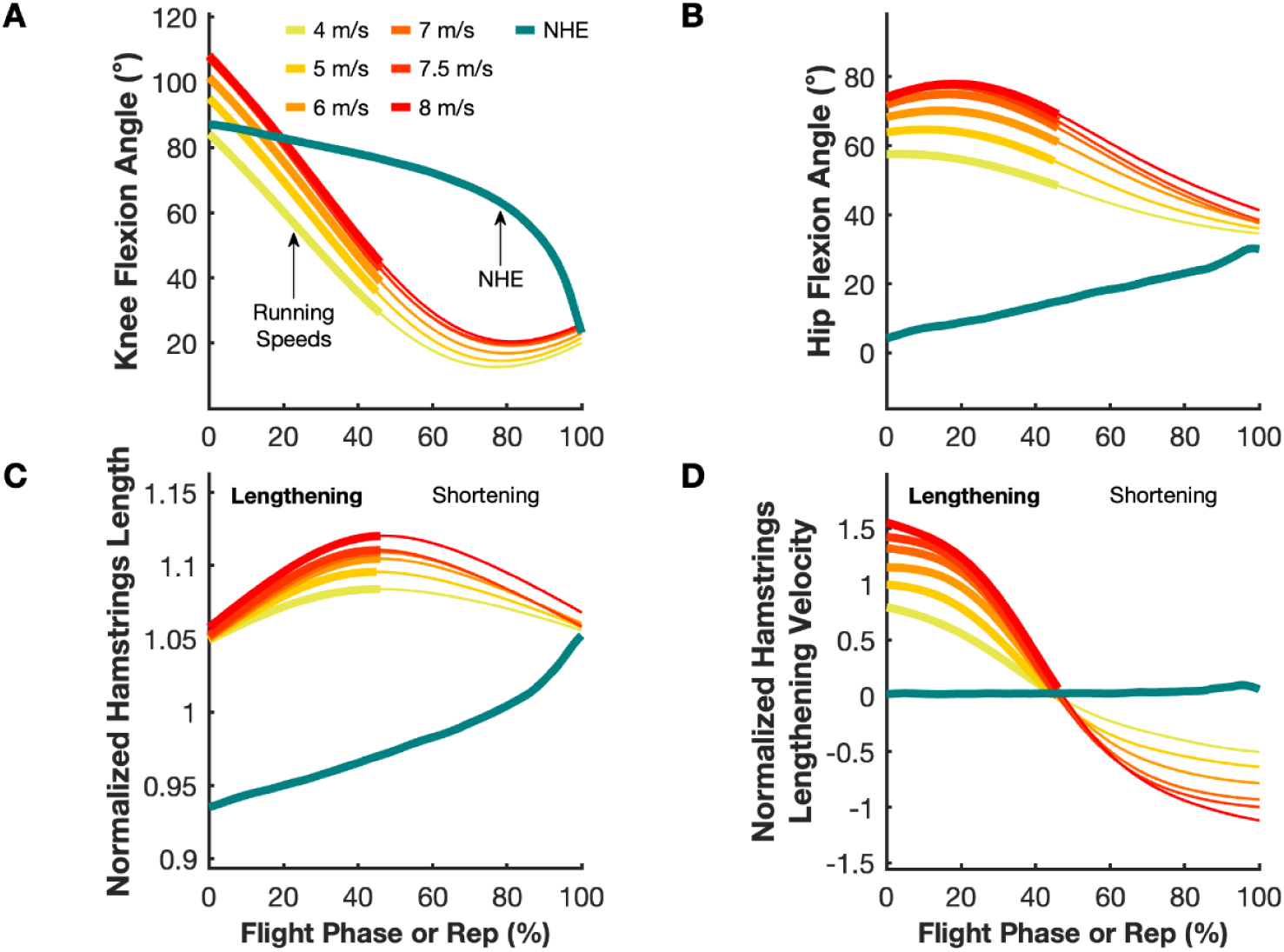
Hip and knee joint and hamstrings kinematics. The curves show mean (A) knee flexion angle, (B) hip flexion angle, (C) normalized hamstrings length, and (D) normalized hamstrings lengthening velocity during the flight phase of running at different speeds and the NHE repetition (Rep). The region over which the hamstrings are lengthening while running is indicated by a thick line; the thinner line is the region over which the hamstrings are shortening. Hamstrings lengths are normalized by the muscle-tendon unit length of the biceps femoris long head with all lower extremity angles set to zero for each subject, and hamstrings velocities are reported in normalized lengths per second.

Peak hamstrings length increased from 1.08 ± 0.02 to 1.12 ± 0.03 as running speed increased from 4 to 8 m/s (*p* < 0.01) and was greater at all speeds compared to the NHE (1.05 ± 0.03) (*p* < 0.001) (Figure 1C). For all running speeds, the hamstrings length initially increased, reaching peak length near the midpoint of the flight phase and then decreased, whereas the hamstrings length increased throughout the NHE, with a faster increase over approximately the last 10% of the repetition. Similarly, peak hamstrings lengthening velocity increased from 0.8 ± 0.09 to 1.55 ± 0.18 normalized lengths/s as running speed increased from 4 to 8 m/s (*p* < 0.05) and was higher at all speeds compared to the NHE (*p* < 0.001) (Figure 1D).

For running, the peak knee flexion moment increased in magnitude from 1.06 ± 0.15 to 2.53 ± 0.50 Nm/kg as speed increased from 4 to 8 m/s (*p* < 0.001) and was greater for all speeds equal to or greater than 6 m/s (*p* < 0.001), when compared to the NHE (Figure 2A). Peak negative knee flexion power increased in magnitude from 11.7 ± 1.5 to 35.5 ± 6.3 W/kg as speed increased from 4 to 7 m/s (*p* < 0.05) but did not continue to increase beyond 7 m/s. Peak negative knee flexion power was greater for all running speeds compared to the NHE (*p* < 0.001) (Figure 2B).

**Figure 2.**
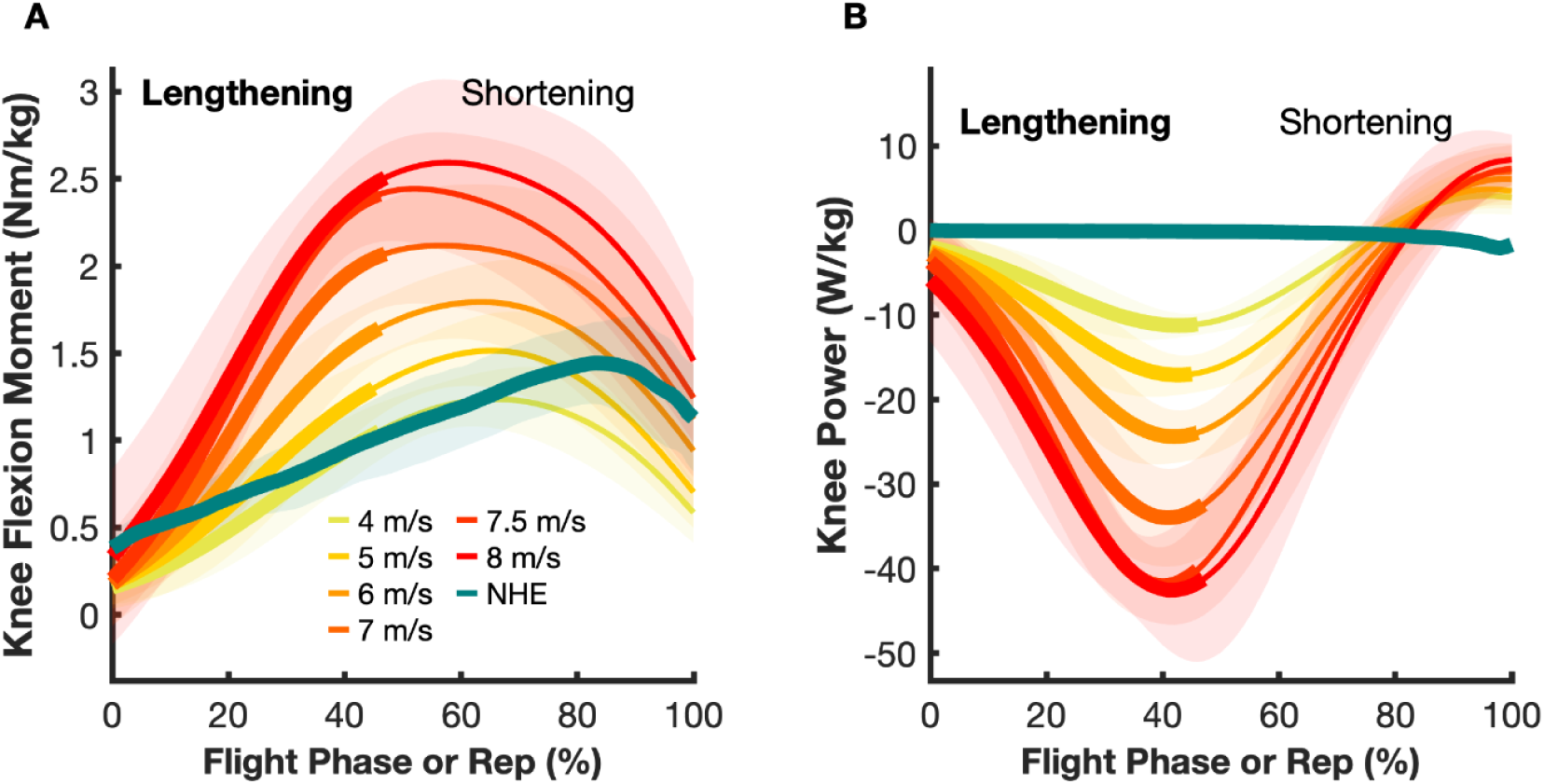
Normalized knee flexion moment (A) and normalized knee power (B) during the flight phase at different running speeds and the NHE (mean ± 1 standard deviation indicated by shaded bands). The hamstrings were lengthening over the region indicated by the thick lines. Note the larger power absorption (negative power) in running compared to NHE. Moments and powers are normalized by body mass.

Peak hip extension moment increased in magnitude from 1.91 ± 0.36 to 4.66 ± 0.96 Nm/kg as running speed increased from 4 to 8 m/s (*p* < 0.05) and was greater at all running speeds compared to the NHE (*p* < 0.001) (Figure 3A). Peak negative hip extension power increased in magnitude from 1.91 ± 0.36 to 8.75 ± 2.85 W/kg as running speed increased from 4 to 8 m/s (*p* < 0.05) and was greater at running speeds of 6 m/s and above compared to the NHE (*p* < 0.001) (Figure 3B).

**Figure 3.**
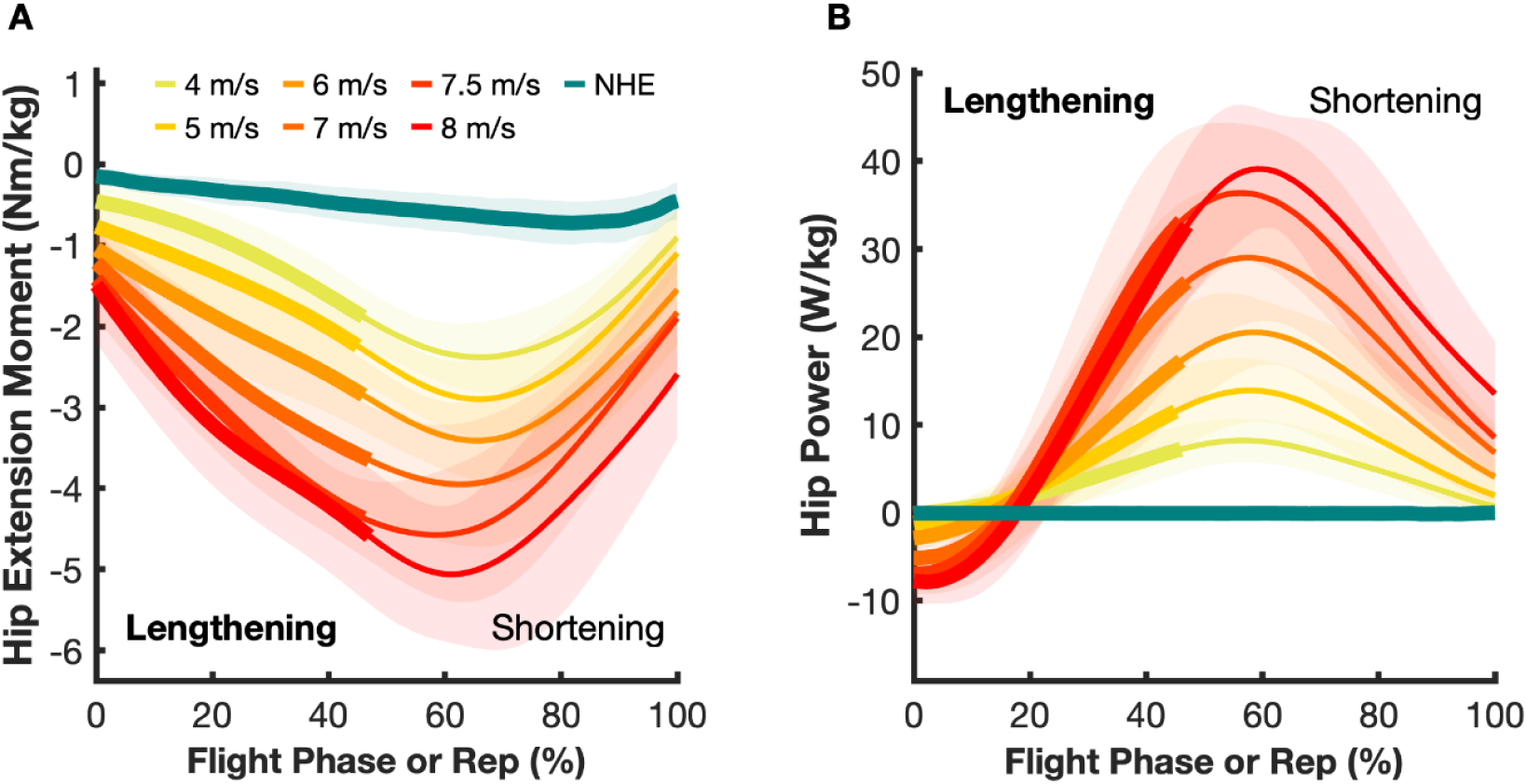
Normalized hip extension moment (A) and normalized hip power (B) during the flight phase of running and the NHE (mean ± 1 standard deviation indicated by shaded bands). The hamstrings were lengthening over the region indicated by the thick lines. Moments and powers are normalized by body mass.

The net knee flexion work was negative when the hamstrings were lengthening in running and during the NHE. For all running speeds and the NHE, the knee flexion moment was always positive, and the angular velocity was always negative while the hamstrings were lengthening. Negative knee flexion work increased in magnitude from 0.50 ± 0.11 to 1.62 ± 0.26 J/kg as speed increased from 4 to 8 m/s (*p* < 0.05). The NHE negative knee flexion work was greater than running at 4-6 m/s (*p* < 0.001); however, running negative knee flexion work was greater than the NHE at 7.5 m/s and above (*p* < 0.001) (Figure 4A).

**Figure 4.**
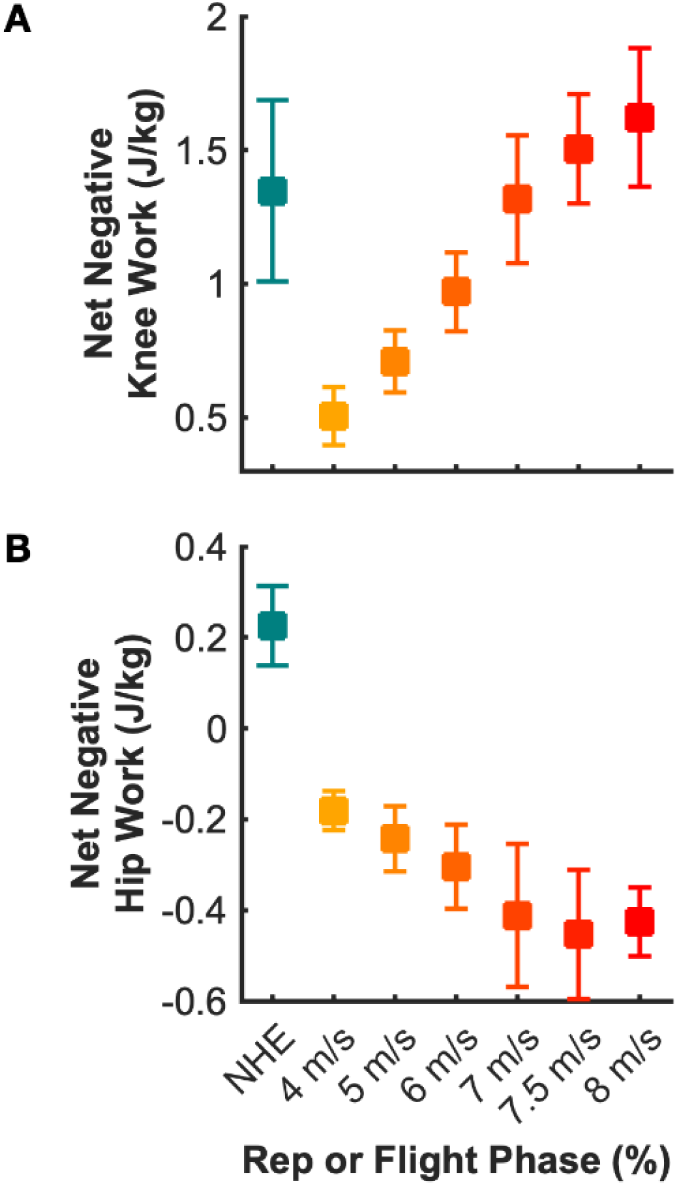
Normalized net negative knee flexion work (A) and normalized net negative hip extension work (B). The values presented are the mean ± 1 standard deviation. The work at the knee and hip were both calculated while the hamstrings are lengthening. Positive values indicate negative work. Work is normalized by body mass.

For all running speeds, the net extension work at the hip was due to a combination of negative and positive hip extension work. The hip extension moment was always negative while the hamstrings were lengthening, but the hip flexion angular velocity polarity changed. For the NHE, the net extension work at the hip was entirely due to negative hip extension work. Net hip extension work was positive for all running speeds, and increased in magnitude from 0.18 ± 0.04 to 0.43 ± 0.08 J/kg as speed increased from 4 to 7 m/s (*p* < 0.05) but did not increase above 7 m/s. Conversely, for the NHE, net hip extension work (−0.23 ± 0.09 J/kg) was negative and differed compared to all running speeds (*p* < 0.001) (Figure 4B).

## 4. Discussion

This study compared the joint-level mechanical loads of high-speed running and the NHE, which revealed that sprinting requires higher knee flexion moments and higher negative knee flexion powers than the NHE. High-speed running also puts the hamstrings at longer lengths and higher lengthening velocities in contrast to the NHE, which is performed at relatively shorter hamstrings lengths and much lower velocities. However, the NHE duration is almost 70 times longer than the hamstrings lengthening portion of the flight phase of running at 8 m/s (Figure 5), indicating that hamstrings force is generated over a much longer time period during the NHE. The net negative knee work during running and the NHE are thus comparable. This characterization of the biomechanical demands of running and the NHE can help guide future exercise studies that examine how the two training modalities affect muscle-tendon adaptation.

**Figure 5.**
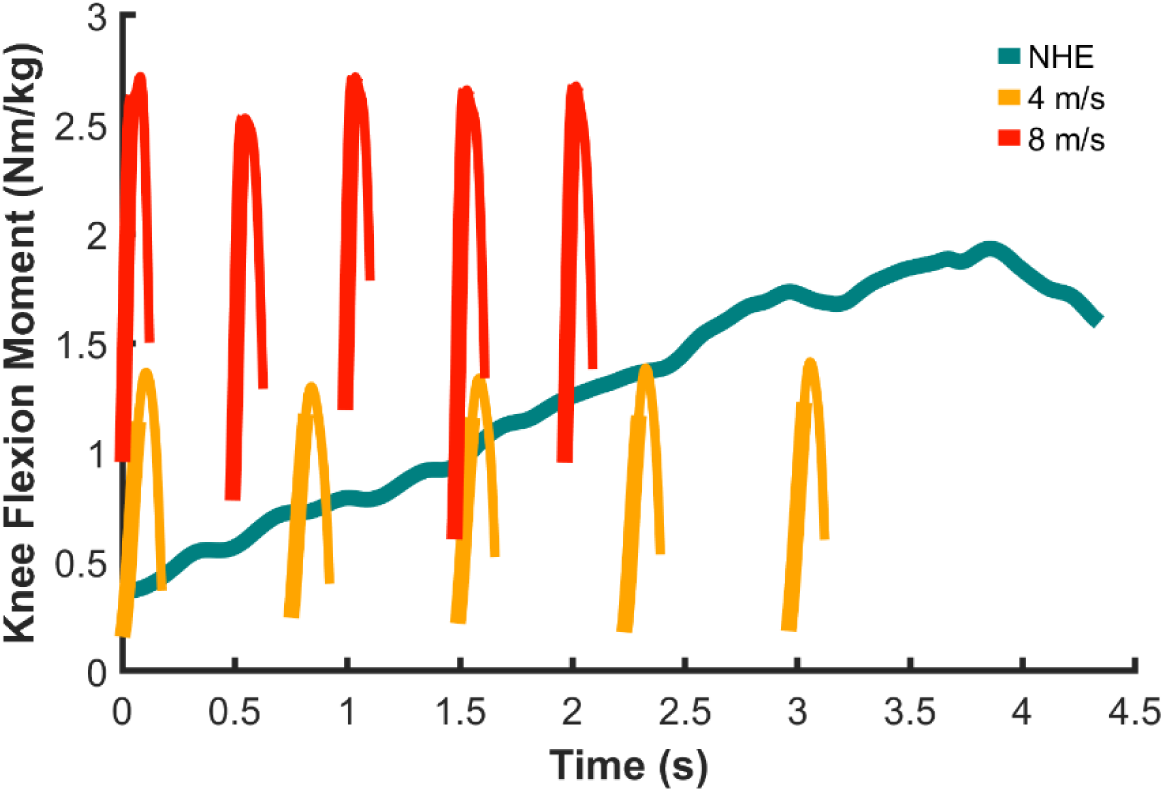
Normalized knee flexion moment during the flight phase of five consecutive strides at 4 and 8 m/s and a single NHE repetition for a representative participant. The hamstrings were lengthening over the region indicated by the thick lines. Moments are normalized by body mass.

Our results are in agreement with prior research that examined the loads of high-speed sprinting or NHEs and extend this past work by quantifying a full suite of joint-level mechanical loads across a range of running speeds and the NHE in the same participants. During the NHE, we found that the hamstrings lengthened slowly until the final phase of the repetition. Similarly, Van Hooren et al.^26^ found the hamstrings lengthened at an increased rate from 88 to 100% of the NHE.^26^ The average peak knee flexion moment reported here (1.52 ± 0.28 Nm/kg) is for one leg, which is slightly less than the peak knee flexion moment measured in a prior NHE training study^19^ (1.88 ± 0.34 Nm/kg for a single leg after 3 weeks of training). The peak knee moments from the flight phase reported by Dorn et al.^37^ were within one to two standard deviations from our peak knee moments from the flight phase of running for matched speeds. Based on these separate, prior studies, NHE peak knee moments would have been about the same as running at approximately 7 m/s, whereas we found that the peak knee flexion moments in running at or above 6 m/s were greater than the NHE.

Our results shed light on potential mechanisms for the effectiveness of the NHE.^17,18^ The NHE requires that athletes generate a substantial knee flexion moment for a much longer time compared to the brief knee flexion moment observed for running. The NHE takes the hamstring muscles to failure because of the high force and long duration, and has been demonstrated in several studies to increase hamstrings fascicle lengths.^19–21^ This difference in mechanical load may be sufficient to lead to hamstring adaptation, given that adaptation can be induced by novel exercises.^41^

On the other hand, the loading differences between the two modalities suggest that running is important for its specificity, transference and functional relevance.^42,43^ We found that running results in greater knee and hip moments and powers and longer hamstrings lengths and lengthening velocities. It has been suggested that for maximal sarcomerogenesis, movements should be completed at high velocity and through the entire joint range of motion,^16^ and observations have shown preliminary evidence of high velocity movements inducing longer fiber lengths.^44–46^ High acceleration training during growth in guinea fowl results in longer optimal fiber lengths^44^ and sprint athletes have longer fascicles in the quadriceps and plantarflexor muscles compared to non-sprinters.^45,46^ The higher MTU lengths and velocities in running may provide stimulus for fascicle lengthening while having high transference to the movement scenario associated with injury. Previous monitoring of sports teams have found that exposure to moderate sprint distances (864-1,453 m) over a three week period during preseason was associated with a significantly lower injury risk than exposure to low (< 864 m) and high (> 1,453 m) sprint distances.^47^ Therefore, moderate amounts of sprint training may partner well with the NHE in injury prevention training programs.

There are a few limitations to consider when interpreting the results of our study. First, some of the participants in our study were not highly trained in the NHE. Despite lower training loads, two of our participants showed similar peak knee moments (1.86 ± 0.21 Nm/kg) to group averages measured immediately after an intensive NHE training intervention study (1.88 ± 0.34 Nm/kg).^19^ Second, our peak knee moments in running (2.12 ± 0.34 Nm/kg at 7.0 m/s) are larger than those reported by Schache et al. (1.08 ± 0.16 Nm/kg at 6.97 ± 0.09 m/s),^36^ but more closely match the peak knee moments reported in Dorn et al. (approximately 1.7 Nm/kg at 7.0 ± 0.1 m/s).^37^ Third, our analysis characterized only two hamstring training modalities, high-speed running and the NHE, whereas other training modalities such as the Romanian deadlift are also used^48,49^ and were not examined in our study. Finally, our conclusions are based on MTU lengths and velocities and joint-level analyses, which do not provide insights into muscle-tendon dynamics or the state of individual muscles. Thus, further work is needed to explore differences in hamstrings fiber lengths and velocities between the two training modalities. While limitations exist with a joint-level analysis, our results provide a baseline understanding for a muscle analysis and are more easily measured by practitioners and in intervention studies.

Taken together, our results show that running and NHEs provide different and complementary biomechanical demands on the muscles crossing the hip and knee. The study’s statistical and biomechanical methods provide a basis for interpreting differences in joint-level mechanics in running and NHEs that can be further enhanced with a muscle-level analysis.

## Acknowledgements

This work was supported by the National Science Foundation, the Joe and Clara Tsai Foundation through the Wu Tsai Human Performance Alliance, and the Stanford Graduate Fellowship in Science and Engineering. We thank Julie Muccini, Julie Kolesar, Tian Tan, and Sam Hamner for their support during the data collection sessions. We also thank Max Andrews and Glen Lichtwark for their assistance in this study, as well as Dawit Lee and Jeff Stribling for their expertise in circuitry and Alex Gonzalez for their help with the statistical analysis.

## Authors’ contributions

KS and NH were responsible for data collection and analysis. All authors participated in the conception and design of the study, the interpretation of results, and manuscript writing. All authors have read and approved the final version of the manuscript and agree with the order of presentation of the authors.

## Data availability statement

The data that support the findings of this study will be made openly available at https://simtk.org/projects/nhe_run_sims.

## References

1. Jones A, Jones G, Greig N, Bower P, Brown J, Hind K. Epidemiology of injury in English Professional Football players: A cohort study. Phys Ther Sport 2019;35:18–22.

2. Ekstrand J, Hägglund M, Kristenson K, Magnusson H, Waldén M. Fewer ligament injuries but no preventive effect on muscle injuries and severe injuries: an 11-year follow-up of the UEFA Champions League injury study. Br J Sports Med 2013;47:732–7.

3. Orchard J, Seward H, Orchard J. Results of 2 decades of injury surveillance and public release of data in the Australian Football League. Am J Sports Med 2013;41:734–41.

4. Malliaropoulos N, Papacostas E, Kiritsi O, Papalada A, Gougoulias N, Maffulli N. Posterior thigh muscle injuries in elite track and field athletes. Am J Sports Med 2010;38:1813–9.

5. Askling C, Tengvar M, Thorstensson A. Acute hamstring injuries in Swedish elite football: A prospective randomised controlled clinical trial comparing two rehabilitation protocols. Br J Sports Med 2013;47:953–9.

6. Ekstrand J, Lee J, Healy J. MRI findings and return to play in football: a prospective analysis of 255 hamstring injuries in the UEFA Elite Club Injury Study. Br J Sports Med 2016;50:738–43.

7. Hägglund M, Waldén M, Magnusson H, Kristenson K, Bengtsson H, Ekstrand J. Injuries affect team performance negatively in professional football: an 11-year follow-up of the UEFA Champions League injury study. Br J Sports Med 2013;47:738–42.

8. Ekstrand J, Waldén M, Hägglund M. Hamstring injuries have increased by 4% annually in men’s professional football, since 2001: a 13-year longitudinal analysis of the UEFA Elite Club injury study. Br J Sports Med 2016;50:731–7.

9. Kenneally-Dabrowski C, Brown N, Lai A, Perriman D, Spratford W, Serpell BG. Late swing or early stance? A narrative review of hamstring injury mechanisms during high-speed running. Scand J Med Sci Sports 2019;29:1083–91.

10. Schache AG, Dorn TW, Blanch PD, Brown NAT, Pandy MG. Mechanics of the human hamstring muscles during sprinting. Med Sci Sports Exerc 2012;44:647–58.

11. Chumanov ES, Heiderscheit BC, Thelen DG. The effect of speed and influence of individual muscles on hamstring mechanics during the swing phase of sprinting. J Biomech 2007;40:3555–62.

12. Chumanov ES, Heiderscheit BC, Thelen DG. Hamstring musculotendon dynamics during stance and swing phases of high speed running. Med Sci Sports Exerc 2011;43:525–32.

13. Yu B, Queen RM, Abbey AN, Liu Y, Moorman CT, Garrett WE. Hamstring muscle kinematics and activation during overground sprinting. J Biomech 2008;41:3121–6.

14. Proske U, Morgan DL, Brockett, Percival P. Identifying athletes at risk of hamstring strains and how to protect them. Clin Exp Pharmacol Physiol 2004;31:546–50.

15. Butterfield TA, Leonard TR, Herzog W. Differential serial sarcomere number adaptations in knee extensor muscles of rats is contraction dependent. J Appl Physiol 2005;99:1352–8.

16. Hinks A, Franchi MV, Power GA. The influence of longitudinal muscle fascicle growth on mechanical function. J Appl Physiol 2022;133:87–103.

17. van der Horst N, Smits D-W, Petersen J, Goedhart EA, Backx FJG. The preventive effect of the Nordic hamstring exercise on hamstring injuries in amateur soccer players: a randomized controlled trial. Am J Sports Med 2015;43:1316–23.

18. Petersen J, Thorborg K, Nielsen MB, Budtz-Jørgensen E, Holmich P. Preventive effect of eccentric training on acute hamstring injuries in men’s soccer: A cluster-randomized controlled trial. Am J Sports Med 2011;39:2296–303.

19. Pincheira PA, Boswell MA, Franchi MV, Delp SL, Lichtwark GA. Biceps femoris long head sarcomere and fascicle length adaptations after 3 weeks of eccentric exercise training. J Sport Health Sci 2022;11:43–9.

20. Ripley NJ, Cuthbert M, Comfort P, McMahon JJ. Effect of additional Nordic hamstring exercise or sprint training on the modifiable risk factors of hamstring strain injuries and performance. PLoS One 2023;18:e0281966.

21. Andrews MH, Pai A, Gurchiek RD, Pincheira PA, Chaudhari AS, Hodges PW, et al. Multiscale hamstring muscle adaptations following 9 weeks of eccentric training. J Sport Health Sci 2024;14:100996.

22. Malone S, Roe M, Doran DA, Gabbett TJ, Collins K. High chronic training loads and exposure to bouts of maximal velocity running reduce injury risk in elite Gaelic football. J Sci Med Sport 2017;20:250–4.

23. Freeman BW, Young WB, Talpey SW, Smyth AM, Pane CL, Carlon TA. The effects of sprint training and the Nordic hamstring exercise on eccentric hamstring strength. J Sports Med Phys Fitness 2019;59:1119–25.

24. van den Tillaar R, Solheim JAB, Bencke J. Comparison of hamstring muscle activation during high-speed running and various hamstring strengthening exercises. Int J Sports Phys Ther 2017;12:718–27.

25. Johnson D, Buckley JG. Muscle power patterns in the mid-acceleration phase of sprinting. J Sports Sci 2001;19:263–72.

26. Van Hooren B, Vanwanseele B, van Rossom S, Teratsias P, Willems P, Drost M, et al. Muscle forces and fascicle behavior during three hamstring exercises. Scand J Med Sci Sports 2022;32:997–1012.

27. Li C, Liu Y. Regional differences in behaviors of fascicle and tendinous tissue of the biceps femoris long head during hamstring exercises. J Electromyogr Kinesiol 2023;72:102812.

28. Ripley N, Fahey J, Comfort P, McMahon J. Kinematic, Neuromuscular and Bicep Femoris In Vivo Mechanics during the Nordic Hamstring Exercise and Variations of the Nordic Hamstring Exercise. Muscles 2024;3:310–22.

29. Raiteri BJ, Beller R, Hahn D. Biceps femoris long head muscle fascicles actively lengthen during the nordic hamstring exercise. Front Sports Act Living 2021;3:669813.

30. Werling K, Bianco NA, Raitor M, Stingel J, Hicks JL, Collins SH, et al. AddBiomechanics: Automating model scaling, inverse kinematics, and inverse dynamics from human motion data through sequential optimization. Plos One 2023;18:e0295152.

31. Lai AK, Arnold AS, Wakeling JM. Why are antagonist muscles co-activated in my simulation? A musculoskeletal model for analysing human locomotor tasks. Ann Biomed Eng 2017;45:2762–74.

32. Dembia CL, Bianco NA, Falisse A, Hicks JL, Delp SL. OpenSim Moco: Musculoskeletal optimal control. PLOS Comput Biol 2020;16:e1008493.

33. Delp SL, Anderson FC, Arnold AS, Loan P, Habib A, John CT, et al. OpenSim: Open-Source Software to Create and Analyze Dynamic Simulations of Movement. IEEE Trans Biomed Eng 2007;54:1940–50.

34. Seth A, Hicks JL, Uchida TK, Habib A, Dembia CL, Dunne JJ, et al. OpenSim: Simulating musculoskeletal dynamics and neuromuscular control to study human and animal movement. PLOS Comput Biol 2018;14:e1006223.

35. Gurchiek RD, Teplin Z, Falisse A, Hicks JL, Delp SL. Hamstrings are stretched more and faster during accelerative running compared to speed-matched constant speed running. Med Sci Sports Exerc 2025;57:461–9.

36. Schache AG, Blanch PD, Dorn TW, Brown NA, Rosemond D, Pandy MG. Effect of running speed on lower limb joint kinetics. Med Sci Sports Exerc 2011;43:1260–71.

37. Dorn TW, Schache AG, Pandy MG. Muscular strategy shift in human running: dependence of running speed on hip and ankle muscle performance. J Exp Biol 2012;215:1944–56.

38. R Core Team. R: A Language and Environment for Statistical Computing [Internet]. Vienna, Austria: R Foundation for Statistical Computing, 2024. Available from: https://www.R-project.org/

39. Bates D, Mächler M, Bolker B, Walker S. Fitting Linear Mixed-Effects Models Using lme4. J Stat Softw 2015;67:1–48.

40. Lenth RV. emmeans: Estimated Marginal Means, aka Least-Squares Means [Internet]. 2025. Available from: https://rvlenth.github.io/emmeans/

41. Zatsiorsky VM, Kraemer WJ, Fry AC. Science and Practice of Strength Training. Human Kinetics, 2021. 346 p.

42. Lee M. Running Biomechanics and Lower Limb Strength Associated with Prior Hamstring Injury. Med Sci Sports Exerc 2009;41:1942–51. 10.1249/MSS.0b013e3181a55200.

43. Vicens-Bordas J, Sarand AP, Beato M, Buhmann R. Hamstring Injuries, From the Clinic to the Field: A Narrative Review Discussing Exercise Transfer. Int J Sports Physiol Perform 2024;19:729–37.

44. Salzano MQ, Suzanne MC, Stephen JP, Rubenson J. High-acceleration training during growth increases optimal muscle fascicle lengths in an avian bipedal model. J Biomech 2018;80:1.

45. Abe T, Fukashiro S, Harada Y, Kawamoto K. Relationship between sprint performance and muscle fascicle length in female sprinters. J Physiol Anthropol Appl Human Sci 2001;20:141–7.

46. Kumagai K, Abe T, Brechue WF, Ryushi T, Takano S, Mizuno M. Sprint performance is related to muscle fascicle length in male 100-m sprinters. J Appl Physiol 2000;88:811–6.

47. Colby MJ, Dawson B, Heasman J, Rogalski B, Gabbett TJ. Accelerometer and GPS-derived running loads and injury risk in elite Australian footballers. J Strength Cond Res 2014;28:2244–52.

